# Animal learning in a multidimensional discrimination task as explained by dimension-specific allocation of attention

**DOI:** 10.1101/271379

**Authors:** Flavia Aluisi, Anna Rubinchik, Genela Morris

**Affiliations:** Sagol Department of Neurobiology; University of Haifa, Abba Khoushy Ave 199, Haifa, 3498838 Israel; Department of Economics; University of Haifa, Abba Khoushy Ave 199, Haifa, 3498838 Israel

**Keywords:** reinforcement learning, attention, set-shifting, rule-learning

## Abstract

Reinforcement learning describes the process by which during a series of trial-and-error attempts, actions that culminate in reward are strengthened. When the actions are based on sensory stimuli, an association is formed between the stimulus, the action and the reward. Computational, behavioral and neurobiological accounts of this process successfully explain simple stimulus-response learning. However, if the cue is multi-dimensional, identifying which of its features are relevant for the reward is not trivial, and the underlying cognitive process is poorly understood. To study this we adapted an intra-dimensional/ extra-dimensional set-shifting paradigm to train rodents on a multidimensional sensory discrimination task. In our setup, stimuli of different modalities (spatial, olfactory and visual) are combined into complex cues and manipulated independently. In each set, only a single stimulus dimension is relevant for reward. To distinguish between learning and decision-making we suggest a weighted attention model (WAM). It combines a learning model where each feature-dimension is reinforced separately with a decision rule that chooses an alternative according to a weighted average of learnt values, in which weight is associated with each dimension. We estimated the parameters of the WAM (decision weights, learning rate and noise) and demonstrated that is outperforms an alternative model in which a value learnt is assigned to each combination of features, or every state. Estimated decision weights of WAM reveal an experience-based bias in learning. The intra-dimensional set shift separated the decision weights. While in the first phase of the experiment the weights were roughly the same, in the second phase the weight on the dimension that was key to finding the reward became higher than others. After the extra-dimensional shift this dimension became irrelevant, however its decision weight remained high for the early learning stage in this last phase, providing an explanation for the poor performance of the animals. By the end of the phase when the rats performance improved, the weights for the two dimensions converged. Thus, estimated weights can be viewed as a possible way to quantify the experience-based bias.

## 1 Introduction

A common conceptualization of choices between alternatives highlights two components to the decision process: outcome valuation and mapping of states and actions to outcomes, (Rangel et al., 2008; Gilboa and Marinacci, 2016). Even if the subjective value of each outcome is known, computing similarity between different states, as required for state-action-outcome mapping, is a hard problem (Erev and Roth, 2014; Argenziano and Gilboa, 2017). Lack of knowledge about the environment might either lead to avoiding of unknown environments, as in Ellsberg (1961), or it may motivate exploration and learning (Rangel et al., 2008). Modeling approaches of the learning process in decision making contexts has differed between fields of research. Whereas in economics it has been predominantly based on the Bayesian framework, (Gilboa and Marinacci, 2016), a frequentist-based reinforcement learning (RL) (Sutton and Barto, 1998) approach was instrumental in explaining human and animal choice behavior and in pointing to the function of the underlying neurobiological structures.

Classical RL models (Wagner and Rescorla, 1972; Sutton and Barto, 1998) assume that a decision-maker assigns a value to each action (or to action-state pair) and updates it according to the reward history. These values form the basis for decisions. However, there is clearly more to adaptive choice behavior than simple updating of values. As similar problems are encountered, learning is signi cantly facilitated. This implies higher level of meta-learning, in which the learning process itself is improved. Meta-learning can take many forms, from tuning of parameters (Doya, 2002) to extracting underlying patterns (Plonsky et al., 2015) or problem characteristics (Collins and Frank, 2013; Gershman and Niv, 2010). Rather than adopting individual solutions, a rule is derived for the family of problems, forming a learning set (Harlow, 1949). Learning sets require uncovering the underlying structure of the task set, allowing generalization. Learning in such scenarios involves two stages: structure learning and parametric learning (Braun et al., 2010).

We will concentrate here on category learning, where categories are naturally de ned by types of sensory input, only a subset of which are relevant to the outcome (Mackintosh and Little, 1969; Roberts et al., 1988). In this common case, it is bene cial to extract only the relevant aspects of the high-dimensional input to simplify learning and decision-making, thus reducing high-dimensional state-spaces to a manageable size. This may be achieved in humans by introducing selective attention to different types of information in the learning process (Slamecka, 1968; Niv et al., 2015), in the decision process, or both (Leong et al., 2017). Rodent studies in which the rules governing reward involve attending to different dimensions of presented stimuli show that, when appropriately trained, the animals’ behavior is consistent with dimension-speci c attention sets (Crofts et al., 2001; Chase et al., 2012; Lindgren et al., 2013; Bissonette and Roesch, 2015; Wright et al., 2015; Aoki et al., 2017). However, it is still not clear how these sets are formed and whether they are learned by the reinforcement schedule. To study how animals deal with extraction of a subset of relevant stimulus dimensions in a high-dimensional setting, we adapted an intra-dimensional/extra-dimensional set shifting paradigm to train rodents on a deterministic multidimensional sensory discrimination task involving spatial, olfactory and visual associations. In each set, only a single stimulus dimension is relevant for reward. To account for the animals’ behavior, we applied a modified reinforcement learning model, combined with a decision rule that chooses among alternatives by comparing the weighted averages of the corresponding learnt values, factored by dimension speci c weights, dubbed *dimension specific attention*. We applied the model to the trial-by-trial choice behavior during the various stages of task performance and show that this model out-performs a simple RL model in describing the data, and that it uncovers patterns of animal behavior that cannot be explained by reinforcement statistics alone.

## 2 Results

### 2.1 Behavior

The data was collected from 18 Long-Evans rats trained on two versions of a multidimensional sensory discrimination task (Fig.1). In this task, naive rats are introduced to a plus-shaped maze. In each trial, the animal is given a choice between two (out of four) randomly chosen arms, indicated by a colored light emitting diodes (LED) and an odor. The pair of odors and the pair of colored LED were randomly assigned to the two doors. For the duration of each phase of training, only one sensory dimension (olfactory, visual or spatial) determined the correct choice. Correct responses were rewarded by a drop of water provided in a port located at the end of the appropriate arm. Training consisted of 50-100 daily trials. We set the threshold of 75% of correct choices during a day as a criterion of successful learning.

**Figure 1:**
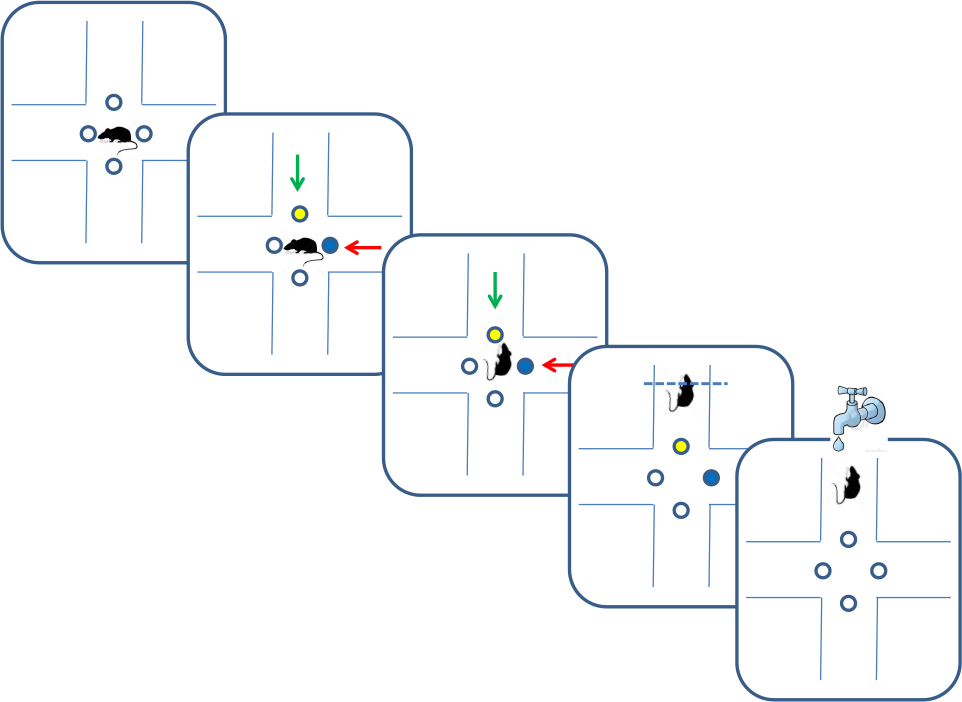
Multidimensional learning paradigm and trial progression. Blue and yellow dots depict illuminated LEDs, and green and red arrows indicate different odors. In the odor phase rats received reward for following the green odor.

Eight animals were trained on an odor-first version of the task and 10 were trained on the LED-first version. In the odor-first group, during the first training phase ODOR 1, the animals reached the 75% success rate within 4.3 ± 0.7 days (*mean ± S.E.M*). In this group, we subsequently performed an intra-dimensional shift, switching the pair of odors used for discrimination. This phase of the task, ODOR 2, proved much easier for the rats, who satis ed the criterion within 1.4 ± 0.3 days, supporting the hypothesis that the animals indeed learned to assign relevance to the correct sensory dimension. Finally we performed an extra-dimensional shift, switching the cue for finding the reward from an odor to a LED color. As expected, the LED phase was substantially more difficult for the animals, requiring 9.2 ± 1.2 days to satisfy the same criterion. Three animals failed to reach the required success rate during this phase (Fig. 2).

**Figure 2:**
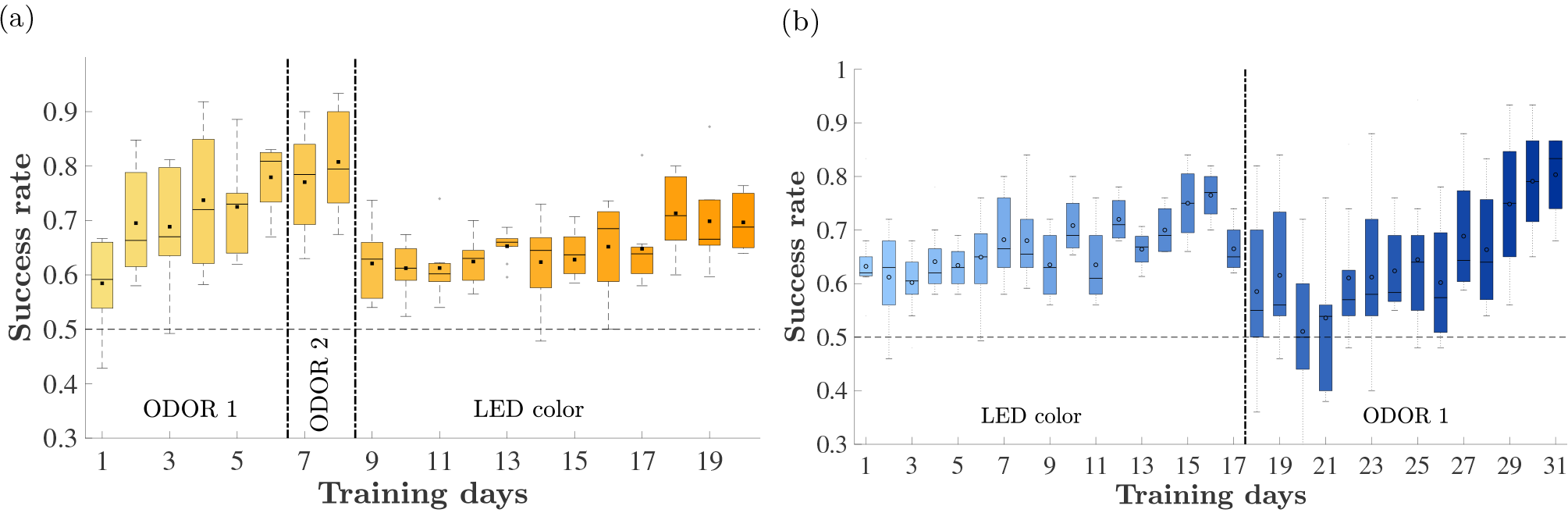
Animals’ performance throughout training: distribution across rats of correct choices as a fraction of total choices by day. Vertical dashed lines depict phase changes in training: the first line depicts the day of the intra-dimensional shift and the second, the day of the inter-dimensional shift. The horizontal dashed line depicts chance level performance. Only days with at least 4 rats training are presented. The black squares depict the mean and the line inside the bar represents the median of the sample, the boundaries of the box denote the 25th and 75th percentile correspondingly. 2a: The odor-first set. 2b: The LED-first set.

The group trained on the LED-first version reached the 75% success rate during the first (LED) training phase within 7.7 ± 1.4 days. After that, the reward rule changed and the relevant stimulus dimension was odor. In this (ODOR 1) phase 7 ± 1.8 days were required to satisfy the successful learning criterion. One rat failed to satisfy the criterion.

### 2.2 Single trial analysis of choice behavior

Trial-to-trial analysis of the animals’ choices allows us to explore learning strategies of the rats and deduce their common features.

For example, a rat may follow a spatial strategy, searching for an arm associated with the highest probability of success. Figure 3 depicts the arms chosen by one rat on four training days of the first phase (ODOR 1). The beginning of training was typically characterized by spatial biases. Figure 3 (top left) depicts the rst day of training in which the animal tended to avoid arm 1 and disproportionately choose arms 2 and 3. This pattern of choice was reinforced by successes associated with arms 2 and 3. The rat’s avoidance of arm 1 also prevented positive reinforcement related to this arm. This behavior appears consistent with learning values assigned to each arm by associating the successes and failures with each location. However, it is not clear whether such value is assigned to every feature (location, type of odor, the color of the LED) independently or to every combination of features.

**Figure 3:**
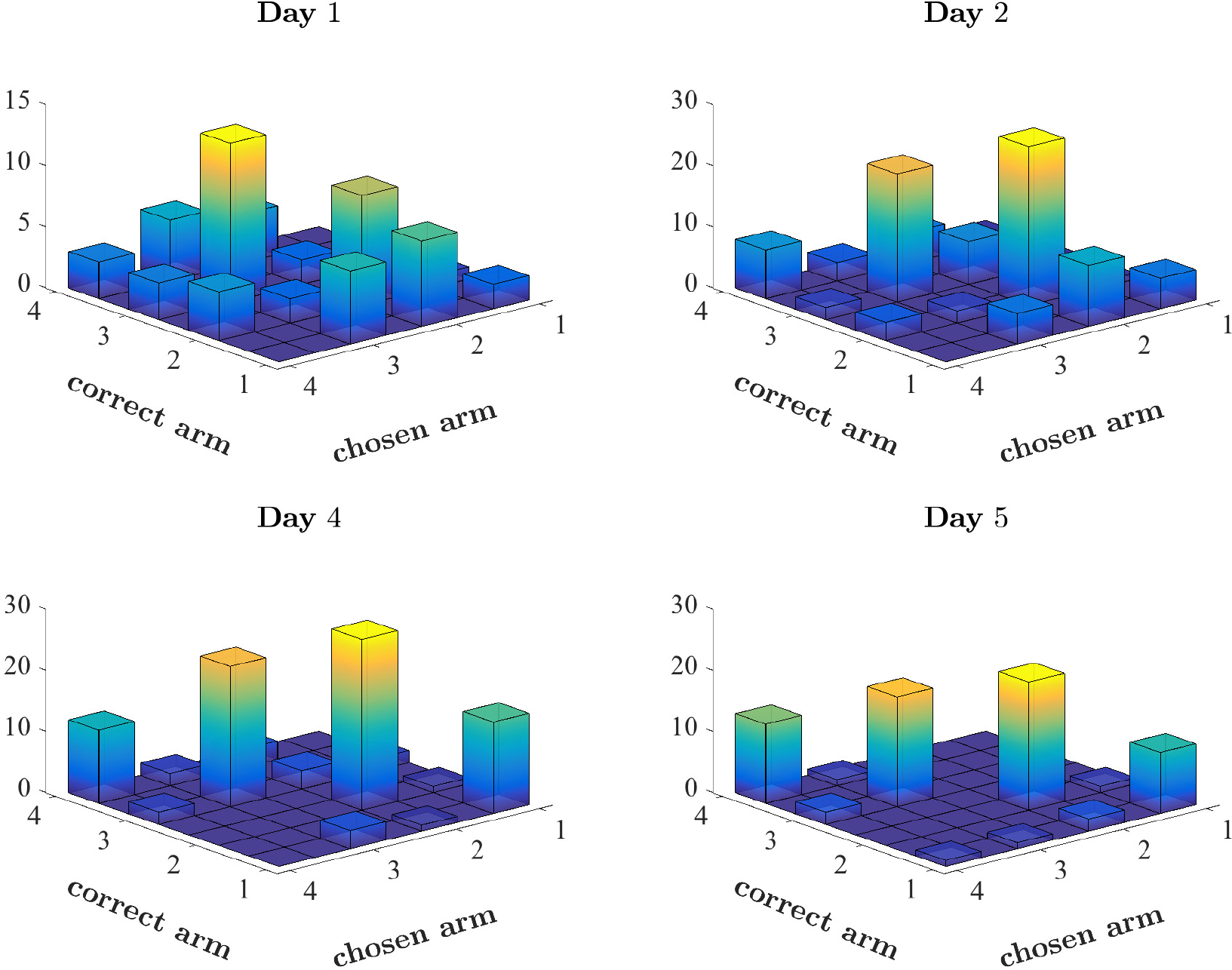
Pattern of choices made by a single rat. Number of times the rat chose each arm plotted against the correct arm in the trial. Correctly performed choices appear on the diagonal. Spatial bias appears as disproportionate number of choices of a single arm (see for example the choices of arm 3 on Day 1). The first two days and the last two days of the first phase of this rat (ODOR 1) are depicted.

Inspection of the later days of the same phase, depicted in the lower panels, reveals that although the bias persists throughout the training, it is less pronounced at the end, leading to a su ciently high rate of success on the last day.

To systematically analyse of the trial-to-trial behavior, we formulate two competing models of learning and decision-making and estimate the parameters of each to fit the actual behavior of the rats. The models are presented formally in Materials and Methods.

The first model, Naive Reinforcement Learning model (NRL, described in section 4.3), is a variation of a classical on-line reinforcement learning procedure (Wagner and Rescorla, 1972; Sutton and Barto, 1998). In this model, every combination of features (e.g., the arm with location 1, apple odor and blue LED) that is associated with a choice has a value that is updated with every choice, i.e., when the rat chooses the arm with that *combination* of features. The model predicts that the alternative with the highest value learned so far will be chosen. However, the decision process is subject to a certain noise level. The model has two parameters, learning and noise. The learning rate is a balancing factor between the effects on the value of the current outcome (success or failure), and the previously acquired value, accumulated before the current trial. Noise is a normally distributed random variable whose standard deviation is a parameter of the model. The noise is added to the value to allow for the classical effect of “exploration” and to account for other factors (other than learning) that a ect the decision.

The Weighted Attention Model (WAM, described in section 4.4), assumes that every feature of the state is reinforced *separately*, i.e., a value is associated with each sensory input. Thus, there is a separate value for each of the four locations 1 – 4, a separate value for each odor (e.g., apple) and, in addition, a pair of values for each of the LED colors. Each of the values of a chosen stimulus is updated after every choice. However, in contrast to the NRL model, the decision is based on a *weighted average of the values* associated with sensory dimensions (location, odor, LED color). These weights are additional parameters of the model, the three weights sum up to one. As in case of the NRL, the WAM also has the learning rate parameter and noise. The parameter estimation procedure for both models is described in detail in section 4.5.

### 2.3 Comparison of the NRL and the WAM in explaining animals’ behavior

We used the two models to explain behavior of 8 rats performing the odor-first version of the task.

We compare the fit of the two models using the standard *overfitting test*. In our case, this test is superior to tests based on log likelihood. Given that the two models are not nested, a standard log likelihood test is meaningless. Moreover, the trials within each training day are not independent, due to the learning dynamics. This makes an application of the Bayesian Information Criterion difficult, as the correct degree of penalty for adding parameters is not easy to determine. We should note, however, that the likelihood generated by the WAM is higher for all animals and phases than that by the NRL.

To perform the test we have estimated parameters of each model on 90% of the trials each day for each rat in every phase and compared the performance of the two models in explaining the behavior in the last 10% of the trials (for every day, rat, and phase). The proportions of trials explained by each model varied between .33 and 1. In 19 out of 27 days, the WAM predicted at least as large a fraction of test trials as the NRL, averaged across rats.

### 2.3.1 The WAM explains mistakes better than the NRL

The NRL underperforms WAM, especially at the initial stages of learning of each new phase, in which WAM captures erroneous choices which NRL cannot. In NRL, the value associated with any stimulus combination that contains the stimulus associated with reward (such as a particular odor in the first phase or a particular color in the third phase) should increase with every correct choice, while all other values should not change and stay equal to the initial value, zero. Hence, if the decision weight on the relevant feature-dimension is not zero, the prediction of the NRL is that the rat will make a correct choice. Figure 4 depicts the fit of both models as a function of the animals’ performance. The fit of the NRL often equals the fraction of correct choices made by a rat, see the concentration of the dots on the diagonal. By contrast WAM explains mistakes, as is evident by the concentration of WAM points above the diagonal.

Mistakes are captured in NRL by adding noise, or “exploration”, i.e., a random deviation from the choice associated with a higher value. However, estimating the magnitude of noise can not help an observer to trace any systematic “pattern of mistakes” or learning dynamics. It is this problem that the WAM is designed to address. Using the WAM we can explore an experience-driven bias in decisions.

**Figure 4:**
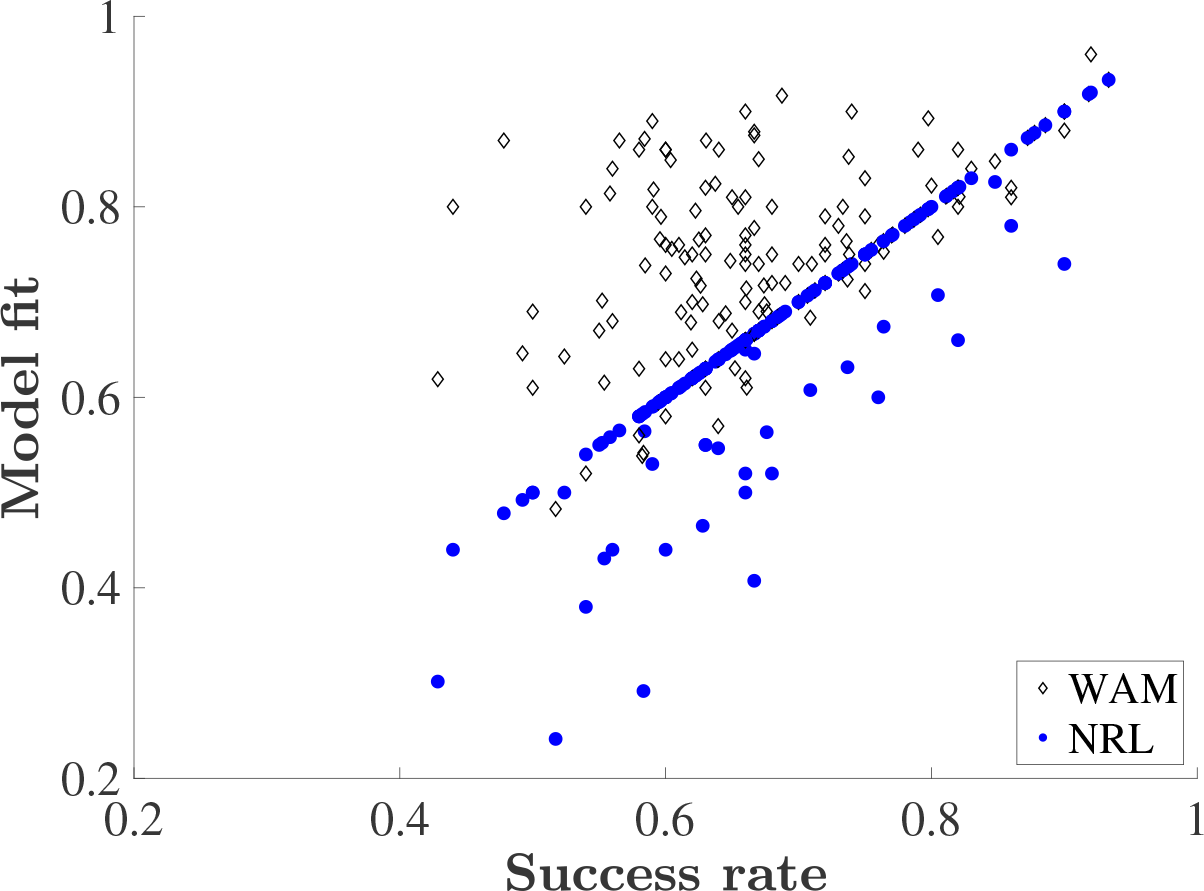
Explanatory power of the WAM versus the NRL. Success rate is the fraction of correct choices made by a rat in a training day. Model fit is the fraction of choices correctly predicted by the corresponding model (WAM and NRL) without adding noise, i.e., based on the accumulated values only.

## 2.4 Exposure to multiple rules induces an exploration over feature-dimension space, causing biased representation

To uncover the learning dynamics we compare the estimated decision weights associated with two different sensory dimensions: olfactory (odor) and visual (LED color). Figure 5 shows the weights associated with the two dimensions throughout the training on the odor-first version of the task. Observation of the weight shift reveals a pattern of an emerging bias. For ODOR 1 phase we find that even as the animals improved, the weights on the relevant feature dimension, odor, did not change signi cantly. However, when an intra-dimensional shift was introduced, an attentional bias appeared, and the weights on odors were higher than those on LED color. After the extra-dimensional shift, the decision weight on odors remained higher than that of the LED colors for the early stage of learning, mirroring the lack of behavioral improvement. As in ODOR 1, even towards the end of this phase, the weights on LED color and odor remained similar.

To examine the temporal changes in estimated decision weights throughout learning, while avoiding biases related to different duration of each phase, we divided each training phase into early and late stages based on each animal’s performance (see Materials and Methods). Figure 6 shows the weights on reward-relevant (left) and reward-irrelevant (right) stimulus dimension in the early and late stages of each phase, averaged by rat. To compare the weights of early and late stages by phase, we used a standard fixed-effects regression explaining the variation in the odor weights. The regression model included a constant term and 2 × 3 stage-phase combinations. Comparisons of the estimated coe cients of each term revealed that weight on odor in the ODOR 1 phase did not change throughout this phase: *P* = 0.89, *F* (1, 35) = 0.018,(Figure 6a). Conversely, the odor weight in the late stage of ODOR 2 is signi cantly different from that in ODOR 1: *P* < 0.05, *F* (1, 35) = 5.41. Interestingly, the odor weight in the late stage of LED did not differ from the early stage of ODOR 1, *P* = 0.99, *F* (1, 35) = 0.0001.

**Figure 5:**
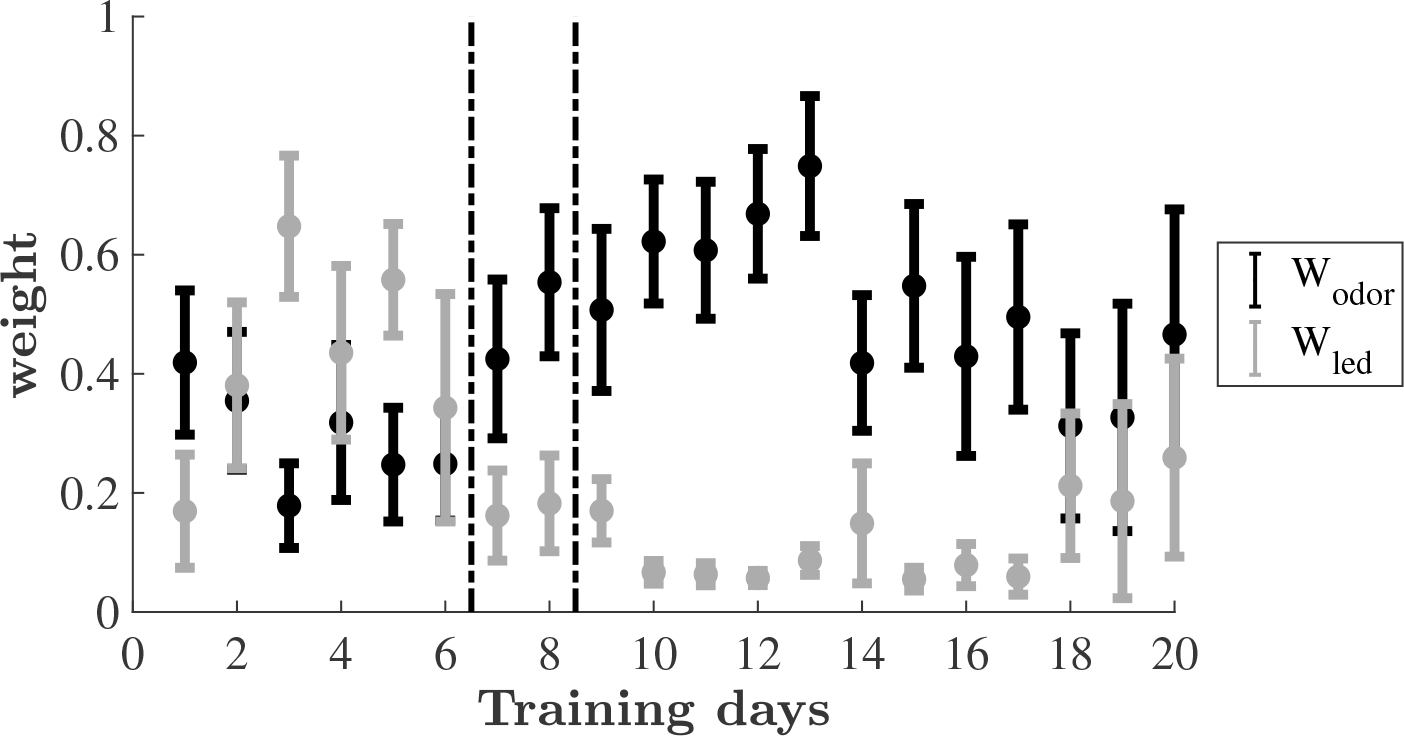
Estimated decision weights on odor and LED color by training days.

**Figure 6:**
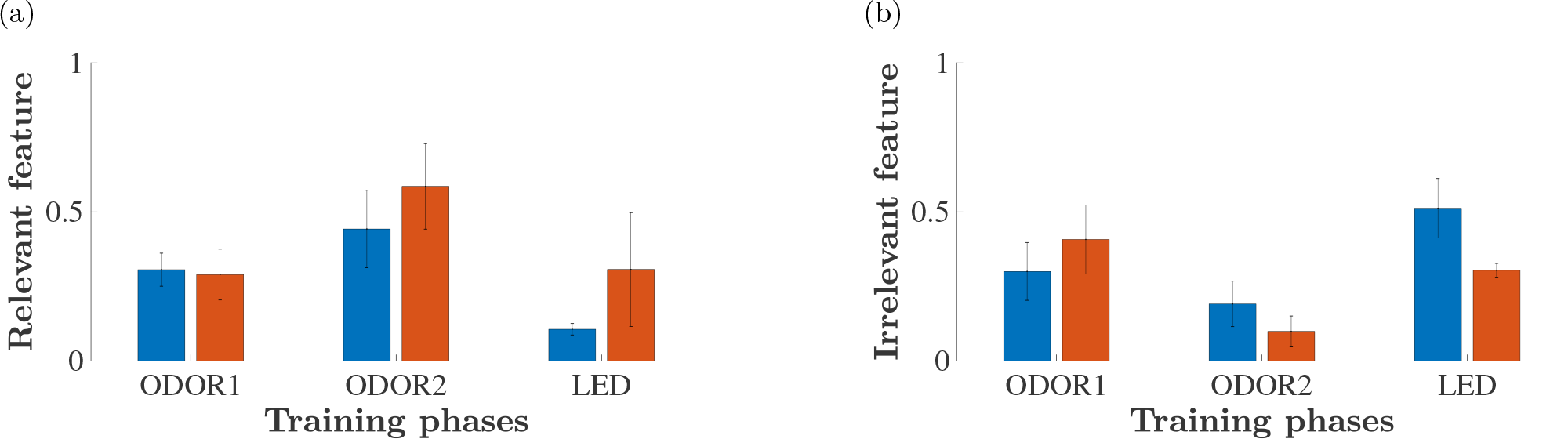
Estimated feature weights. 6a. Weight on relevant feature pooled across animals and days in each phase. 6b. Weight on irrelevant feature pooled across animals and days in each phase in the early (blue) and late (red) stage of learning. LED color is irrelevant in the first two phases, odor is irrelevant in the last phase.

Next, we compared the odor and LED weights throughout the training. We used a fixed effects regression of weights, which has two dimensions: ODOR and LED color, on stage-phase-dimension (2 × 3 × 2) combinations. We found that in ODOR 2 the weights on LED and ODOR in the late stage are signi cantly different: *P* < 0.001, *F* (1, 70) = 14.31. Likewise in LED phase in the early stage the weight on LED and ODOR are significantly different: *P* < 0.05, *F* (1, 70) = 11.37. However, for the late stage of LED the weights on odor and LED are no longer different: *P* = 0.99, *F* (1, 70) = 0.0001.

## 3 Discussion

Animals and humans are commonly confronted with complex situations, in which the most appropriate response is determined by signals that are often of a multidimensional nature. Identifying key features determining payoff, or even the important category (dimension) in each situation relies both on experience and on the underlying model of the world. In the series of experiments described here the dimensions are well de ned: location, odor and LED color. Moreover, the reward schedule is deterministic. Therefore the task of the learner is two-fold: identify the relevant dimension and find the feature associated with reward within this dimension. The weighted reinforcement model introduced here allows to track the values associated with each feature independently of others and then use the weighted average of these values to form a decision. Trial by trial behavior analysis of behavior in a multidimensional task offers several insights. In particular, examining the pattern of incorrect choices made by the animals may uncover the underlying nature of their learning and decision process. The WAM that we use separates learning from decision-making: learning is driven by strengthening or weakening connections between the stimuli and reward, whereas decision-making is based on a weighted sum of learned associations. This dissociation implies two different sources of mistakes. Erroneous choices can be due to unsuccessful learning. Alternatively, it may be that the decision process is faulty, assigning incorrect weights to different feature-dimensions of the problem. Indeed, by separating these two behavioral components, the WAM outperformed the NRL model in explaining the trial-by-trial behavior.

In the final learning phase, after the extra-dimensional shift, the weights associated with the relevant feature were consistently lower than in previous phases, across animals. Inspection of the dynamics of weights throughout learning revealed that for most animals the first days of training on the LEDs phase was dominated by a high weight on the previously relevant feature-dimension (odor). This weight was reflected in the poor performance and slow improvement displayed by these animals. Thus, this result may hint at a mechanism underlying the Einstellung effect (Luchins and Luchins, 1959), in which humans after learning a task exhibit a long latent period before successfully learning a second one.

In a recent paper, Leong et al. find that throughout learning attention is biased towards the relevant category (Leong et al., 2017). Similarly, the decision making model that we use involves comparing weighted scores of available options, where each dimension of a problem is associated with its weight, which in turn, is a-priori independent of the value learned for different features. Thus we, too, separate the process of associating features with outcome (reward) from the process of deciding which aspects (dimensions) of the problem to take into account when comparing available options. However, we do not use observable data to measure the weights, interpreted as allocation of attention, as in Leong et al. (2017), rather, we use the model to identify the weights given the behavioral data only. In other words, our weights are the best ones that explain the behavior, within the model we suggest. The advantage of our approach is that we do not need to rely on a particular observable data. However, we cannot estimate the weights trial-by-trial, instead we fit them day-by-day (for each animal). Despite the variability of the estimates, some visible patterns emerged, as discussed above, and those are quite different from Leong et al. (2017).

In summary, the superior performance of the WAM in explaining trial-by-trial behavior seems to point to a general principle of reinforcement learning in multidimensional environments: rather than associating a state (value) with a combination of features, learning may occur separately along several dimensions. The caveat however is that such conclusion could be a result of our task design. Whether the same principle holds for learning other classes of tasks is left for future research.

## 4 Material and Methods

### 4.1 Animals

18 male Long-Evans rats 8-10 weeks old at the beginning of the experiment were used in this study. The rats were maintained in an inverted 12 h light/12 h dark cycle with unlimited access to food. During training days water consumption was restricted to 1/2 hour every 24 hours. Animal weight and wellbeing were monitored continuously. All experimental procedures were conducted in strict accordance with Institutional Animal Care and Use Committee of the Haifa University (*Ethics approval No 334/14*), as well as the EU and NIH rules and regulations for the use of animals in science research. The animals underwent surgery for implantation of tetrodes for electrophysiological recordings either in their dorsomedial or dorsolateral striatum or in hippocampal area CA1. Results from these recordings are not reported here.

### 4.2 Apparatus and behavioral task

The apparatus used for these experimental tasks was a black plexiglass plus-shaped maze with four 10 x 60 cm arms and a 30×30 cm central hub, situated 1 m above the floor. The entrances to the arms were blocked by retractable automatic doors. Infra-red (IR) sensors at both ends of the arm marked times of arm entry and track completion. The central arm was equipped with 4 nose-poke apparati located on the floor 3 cm away from each door. The nose poke devices had IR sensors to detect nose entry, 3 colored LEDs (blue, yellow and green), and two tubes through which odorised air was delivered and pumped out. Each odor delivery tube could deliver 2 different odors. The odors used were commercial odors that are regularly used in the cosmetics and food industry, diluted 1:1000 from commercial concentrated liquid. Each arm was marked with different large visible cues along the walls. Experiments were conducted in dim light. Each trial started with the rats located in the central hub of the maze. After a variable inter-trial-interval, a buzz was sounded and 2 pseudo-randomly chosen nose pokes were illuminated by LEDs of different colors. Two different odors were delivered to the same nose-pokes and pumped out to keep the odors localized to the nose-poke device. The animals initiated a trial by poking their noses into one of the two nose pokes (or both). Upon this poking, the doors of the corresponding two arms opened and allowed access to the arms. This allowed the rats to choose an arm. If the correct arm was chosen and traversed within the allowed maximum time of 5 seconds, the rats were rewarded by 0.3 ml of water delivered at a reward port at the end of the arm. The correct arm depended on the experimental condition. In the odor phase of the experiment, reward was delivered in the arm associated with a speci c odor (and a random LED) and for the LED condition, reward was delivered in the arm associated with a specific LED (and a random odor). Rats were trained for 50-100 daily trials until they satis ed the criterion performance of 75% correctly performed trials. Following the first odor phase, a different pair of odors was used and the animals had to associate the new odors with the reward. When animals reached the threshold performance for this pair as well they underwent a rule shift and had to learn to associate the reward with one of the two possible LED colors.

### 4.3 Naive reinforcement learning model

#### 4.3.1 Values

As a first approximation, we look for the best fit of the “naïve reinforcement learning” model (NRL) (Niv et al., 2015) to the behavioral data. Every combination of observable features (e.g., arm 3, green LED and apple odor) is de ned as a “state” whose value is to be learned. Thus, if at trial *t* an arm *i* with odor *j* and LED color *k* is chosen, the corresponding value *V*_*i,j,k, t*_ is updated. Update follows the reinforcement learning (RL) rule,

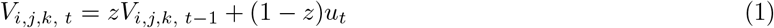

where *z* ∈ [0, 1] is a memory strength parameter, *u*_*t*_ = 1 if the reward was received and *u*_*t*_ = 0 otherwise. The memory parameter *z* used here is equivalent to 1 − *α*, where *α* is the learning rate in the classical reinforcement learning models, cf. e.g. (Sutton and Barto, 1998). Initial values *V*_*i,j,k, t*=0_ are set to zero. Values corresponding to non-chosen options are not updated. Note that the values V are between zero and one. The memory parameter *z* is estimated from the behavioral data separately, as explained in Section *Model Estimation Procedure*.

#### 4.3.2 Decision Rule

Decisions are based on the values associated with each available option. Initial values are set equal to zero. At every trial *t* we calculate the difference in learned values Δ*V*_*t*_ between the values of the two arms that the animal can choose from: a correct and an incorrect one. A deterministic decision rule states that the predicted choice will follow the option with the higher value. Thus, when Δ*V*_*t*_ > 0, the deterministic NRL model predicts that the rat will choose correctly. Since the value of combinations that contain the rewarded feature approach 1 and all other values approach zero, it follows that asymptotically (as *t* → ∞), Δ*V*_*t*_ monotonically converges to 1. Notice that for *z* < 1, after a single success, the value of the chosen action becomes positive. Therefore, starting from the next time the same combination is encountered, the prediction is that the same choice will be repeated. Thus, it is not surprising that the proportion of correct predictions generated by this model, or the *deterministic fit*, should, after all combinations have been encountered at least once, coincide with the success rate: only successes can be predicted but not erroneous choices, cf. fig. 4.

To account for a stochastic nature of animals’ choices, we assume that all other factors a ecting the choice are uncorrelated with the stimulus features and model their effect as a normally distributed noise *ε*^*σ*^ with standard deviation *σ*. This noise may also be interpreted as *exploration*, determining the probability of choosing an action which does not have the highest value learned so far. Similar to *z*, *σ* is estimated from behavior for each day, as explained in Section *Model Estimation Procedure*. A stochastic model predicts that a correct arm will be chosen if the value of a random normal variable with mean Δ*V*_*t*_ and standard deviation is positive. Note that Δ*V*_*t*_ ≥ 0, so the probability is always at least 0.5. As Δ*V*_*t*_ for every pair of combinations that can appear simultaneously at any trial increases over time, so is the predicted probability of choosing the rewarded arm. The rate at which this probability grows depends on *z*. With higher Δ*V*_*t*_ the noise parameter should be higher in order to fit the data with the same proportion of mistakes. To capture the dynamic nature of the learning process, we allow the free parameters to change daily. Parameter estimation was performed by log-likelihood maximization. Updated values for each feature combination were carried over between consecutive days of each learning phase (see detailed description in section *Model estimation procedure*).

### 4.4 The weighted attention model

To capture the animals’ learning and choice behavior better, we devised a modified reinforcement learning model which incorporates feature-by-feature reinforcement with a decision process that employs differential attention to distinct feature dimensions, the weighted attention model (WAM).

The model generates a prediction of the choices made by a rat in each trial. The two basic components of the model are the *values*, which are computed separately for each alternative, stored in memory and updated after each trial according to the *learning rule* below, see eq. (2)-(4), and the *decision rule*, see eq. (5).

#### 4.4.1 The values

In any given trial, action values have 3 components reflecting the three feature dimensions: location, odor and LED color. By design, only one of these components was relevant for getting the reward. Location was never relevant in these experiments (unbeknownst to the rats).

The location component contains 4 variables indicating the values for each of the four arms, cf. Figure 1.
Denote the values at trial *t* by (*l*_1*t*_, *l*_2*t*_, *l*_3*t*_, *l*_4*t*_).

The odor component is a pair, containing the values for each of the two distinct odors: (*o*_1*t*_, *o*_2*t*_). In the odor phase, we let *o*_1*t*_ be the value corresponding to the correct odor, which is the key to finding the reward, whereas *o*_2*t*_ be the value of the second odor, incorrect odor, never associated with the reward.

The LED color component at trial *t* is also a pair, (*c*_1*t*_, *c*_2*t*_). Likewise, in the LED phase the first value is associated with the *correct* LED color.

At the beginning of the series of experiments (*t* = 0) all the values are set to zero.

Values are updated on each trial *t* after the rat has completed its choice and received the feedback (reward/no reward). Assume at trial *t* the animal chose location *i* with odor *j* and LED color *k*. Then the following three variables are updated according to the same reinforcement learning (RL) rule: 
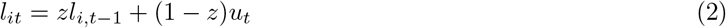

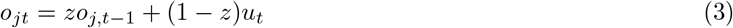

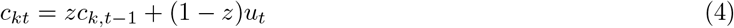
where *z* ∈ [0, 1] is the memory parameter, and *u*_*t*_ is the reward in trial *t*, which is 1 in case of success and 0 in case of failure. The values of all the unchosen features in a given trial are not updated.

#### 4.4.2 The decision rule

The decision rule can be a ected by the values corresponding to the three observable dimensions of each available alternative: location, odor and color. Each value a ects the decision through a corresponding relative weight, *w*_*l*_, *w*_*o*_, *w*_*c*_. All the weights are positive and sum up to 1.

For example, a choice of arm *i*, with odor *j* and LED color *k* receives the composite value score 
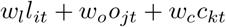

Choices are made by comparison of the composite value scores, and their difference is the decision index. In our example, if arm *i* is the correct arm with odor *j* and LED color *k*, and the other available arm is *i′* with odor *j′* and LED color *k′*, then the decision index is 
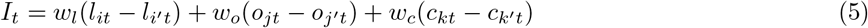

The deterministic model predicts a correct choice if *I*_*t*_ > 0 and an incorrect choice otherwise.

As in the naive reinforcement learning model, we add a stochastic element to this decision rule, so that the predicted probability of a wrong choice is normally distributed with mean *I*_*t*_ and standard deviation *σ*.

### 4.5 Model estimation procedure

The parameters of the NRL model, the memory parameter and the noise, {*z*_*i*_, *σ*_*i*_}, for every day *i* of a given phase, were estimated separately for each phase of the experiment. The memory parameter is restricted to be between zero and unity as it is unity minus the learning rate which we assume to be positive, so 0 ≤ *z*_*i*_ ≤ 1. The noise parameter is non-negative, *σ*_*i*_ > 0 and to keep the speci cation meaningful we restricted it to be less than 30: for higher values the model’s predictions are close to those of a coin toss.

Within every learning phase the values for each combination of features are updated trial-by-trial (according to eq. (1)) and carried over to the following day. These values are set to zero at the beginning of each phase. Parameter estimation is performed by log-likelihood maximization. The maximization was implemented using a built-in constrained optimization routine in Matlab with the initial guess being the best parameter combination found using the simple search over a thin grid of possible parameters.

The parameters of the WAM are the three weights, the memory parameter and the noise for every day *i* of a given phase: {*w*_*i*_ = (*w*_*li*_, *w*_*oi*_, *w*_*ci*_), *z*_*i*_, *σ*_*i*_}, where the three weights 0 ≤ *w*_*li*_, *w*_*oi*_, *w*_*ci*_ ≤ 1 sum to 1. The rest of the parameters satisfy the same constraints as under the NRL.

The parameters are estimated separately for every phase of the experiment, as for the previous model. Within every learning phase the values associated with each feature are updated trial-by-trial (according to eq. 2-4) and carried over to the following day. These values are set to zero at the beginning of each phase. The estimation procedure otherwise is similar to that of the NRL model.

Parameter estimates in both cases are robust to small decay between days, i.e., carrying over to the next day a discounted array of values.

### 4.6 Statistical analysis

To compare the weights on the relevant and irrelevant dimensions in each of the phases we divided each learning phase into an early and late stage according to animal performance. We determined a threshold of 70% for each animal as the first day of the late stage. For each of the 8 animals in the odor-first set and 3 phases we calculated the average weight over the training days of each of the 2 stages.

The weights in each stage and phase were compared by fitting a linear mixed-effects model. Weights on the odor across phases were compared using stage and phase fixed effects. Weights on odor versus LED were compared using dimension, stage and phase fixed effects.

## Author Contributions

GM designed the experiments, FA performed the experiments, AR designed the model, FA,AR and GM analyzed the data, AR and GM wrote the manuscript.

## Acknowledgments

We are grateful to Ediff Barkai for extended discussions and inspiration and George Kour and Avi Mendelsohn for critical reading and helpful discussions.

## References

Aoki, S., Liu, A. W., Zucca, A., Zucca, S., and Wickens, J. R. (2017). New variations for strategy set-shifting in the rat. JoVE (Journal of Visualized Experiments), e55005–e55005

Argenziano, R. and Gilboa, I. (2017). Second-order induction and the importance of precedents. Mimeo, Tel-Aviv University

Bissonette, G. B. and Roesch, M. R. (2015). Rule encoding in dorsal striatum impacts action selection. European Journal of Neuroscience 42, 2555–2567

Braun, D. A., Mehring, C., and Wolpert, D. M. (2010). Structure learning in action. Behavioural brain research 206, 157–165

Chase, E. A., Tait, D. S., and Brown, V. J. (2012). Lesions of the orbital prefrontal cortex impair the formation of attentional set in rats. European Journal of Neuroscience 36, 2368–2375

Collins, A. G. and Frank, M. J. (2013). Cognitive control over learning: creating, clustering, and generalizing task-set structure. Psychological review 120, 190

Crofts, H., Dalley, J., Collins, P., Van Denderen, J., Everitt, B., Robbins, T., et al. (2001). Differential effects of 6-ohda lesions of the frontal cortex and caudate nucleus on the ability to acquire an attentional set. Cerebral Cortex 11, 1015–1026

Doya, K. (2002). Metalearning and neuromodulation. Neural Networks 15, 495–506

Ellsberg, D. (1961). Risk, ambiguity, and the savage axioms. The quarterly journal of economics, 643–669

Erev, I. and Roth, A. E. (2014). Maximization, learning, and economic behavior. Proceedings of the National Academy of Sciences 111, 10818–10825

Gershman, S. J. and Niv, Y. (2010). Learning latent structure: carving nature at its joints. Current opinion in neurobiology 20, 251–256

Gilboa, I. and Marinacci, M. (2016). Ambiguity and the bayesian paradigm. In Readings in Formal Epistemology (Springer). 385–439

Harlow, H. F. (1949). The formation of learning sets. Psychological review 56, 51

Leong, Y. C., Radulescu, A., Daniel, R., DeWoskin, V., and Niv, Y. (2017). Dynamic interaction between reinforcement learning and attention in multidimensional environments. Neuron 93, 451–463

Lindgren, H. S., Wickens, R., Tait, D. S., Brown, V. J., and Dunnett, S. B. (2013). Lesions of the dorsomedial striatum impair formation of attentional set in rats. Neuropharmacology 71, 148–153

Luchins, A. S. and Luchins, E. H. (1959). Rigidity of behavior: A variational approach to the effect of Einstellung. (University of Oregon Press)

Mackintosh, N. and Little, L. (1969). Intradimensional and extradimensional shift learning by pigeons. Psychonomic Science 14, 5–6

Niv, Y., Daniel, R., Geana, A., Gershman, S. J., Leong, Y. C., Radulescu, A., et al. (2015). Reinforcement learning in multidimensional environments relies on attention mechanisms. Journal of Neuroscience 35, 8145–8157

Plonsky, O., Teodorescu, K., and Erev, I. (2015). Reliance on small samples, the wavy recency effect, and similarity-based learning. Psychological review 122, 621–47

Rangel, A., Camerer, C., and Montague, P. R. (2008). A framework for studying the neurobiology of value-based decision making. Nat Rev Neurosci 9, 545–556

Roberts, A., Robbins, T., and Everitt, B. (1988). The effects of intradimensional and extradimensional shifts on visual discrimination learning in humans and non-human primates. The Quarterly Journal of Experimental Psychology 40, 321–341

Slamecka, N. J. (1968). A methodological analysis of shift paradigms in human discrimination learning. Psychological Bulletin 69, 423

Sutton, R. S. and Barto, A. G. (1998). Reinforcement learning: An introduction (Cambridge: MIT Press)

Wagner, A. and Rescorla, R. (1972). Inhibition in pavlovian conditioning: Application of a theory. In Inhibition and learning, eds. R. A. Boakes and M. S. Halliday (London: Academic Press). 301–336

Wright, N. F., Vann, S. D., Aggleton, J. P., and Nelson, A. J. (2015). A critical role for the anterior thalamus in directing attention to task-relevant stimuli. Journal of Neuroscience 35, 5480–5488

